# Posteruptive Loss of Enamel Proteins Concurs with Gain in Enamel Hardness

**DOI:** 10.1101/2024.05.23.595034

**Authors:** Hakan Karaaslan, Alejandro R. Walker, Ana Gil-Bona, Baptiste Depalle, Felicitas B. Bidlack

## Abstract

Tooth enamel maturation requires the removal of proteins from the mineralizing enamel matrix to allow for crystallite growth until full hardness is reached to meet the mechanical needs of mastication. While this process takes up to several years in humans before the tooth erupts, it is greatly accelerated in in the faster developing pig. As a result, pig teeth erupt with softer, protein-rich enamel that is similar to hypomineralized human enamel but continues to harden quickly after eruption.Proteins, such as albumin, that bind to enamel crystals and prevent crystal growth and enamel hardening have been suggested as cause for hypomineralized human enamel that does not naturally harden after eruption. However, albumin is abundant in pig enamel. It is unclear whether fast posteruptive enamel hardening in pigs occurs despite the high protein content or requires a facilitated protein loss to allow for crystal growth.

This study asked how the protein content in porcine enamel changes after eruption in relation to saliva. Based on previous data demonstrating the high albumin content in erupted porcine enamel, we hypothesize that following pre-eruptive maturation, enamel and saliva derived enzymes facilitate protein removal from porcine enamel after eruption. We analyzed enamel and the saliva proteome at three critical timepoints: at the time of tooth eruption, 2 weeks after eruption, and enamel 6 weeks after eruption. We used only fourth deciduous premolars and saliva samples from animals sacrificed at the respective time points to determine the organic content in tooth enamel, saliva, and saliva proteins within enamel.

We found a decrease in the number of proteins and their abundancy in enamel with posteruptive time, including a decrease in serum albumin within enamel. The rapid decrease in the first two weeks is in line with previously reported rapid increase in mineral density of porcine enamel after eruption. In addition to the enamel proteases KLK-4 and MMP-20, we identified serine-, cysteine-, aspartic-, and metalloproteases. Some of these were only identified in enamel, while almost half of the enzymes are in common with saliva at all timepoints. Our findings suggest that the fast posteruptive enamel maturation in the porcine model coincides with saliva exchange and influx of saliva enzymes into porous enamel.

## INTRODUCTION

Healthy tooth enamel forms in a mineralizing matrix that is first secreted until the full enamel thickness is reached. Then follows the cleavage by enamel proteases and extensive removal of matrix proteins to allow for the controlled growth and maturation of crystallites before tooth eruption. This process is slow, occurs before tooth eruption, and provides enamel with its high mineral density and extraordinary hardness, with only 1% organic content by weight remaining in erupted, healthy enamel in adult teeth (Deakins and Volker 1941; Gil-Bona and Bidlack 2020). In contrast, pig teeth form four times faster producing enamel that is much softer compared to healthy human enamel at the time of eruption (Tonge and McCance 1973; Bivin and Mc Clure 1976). At this stage, the hardness and the protein composition of pig enamel resembles hypomineralized enamel in the demarcated opacities seen in the human developmental dental defect of Molar Hypomineralization (MH) (Farah et al. 2010; Gil-Bona et al. 2023). In contrast to MH enamel lesions that are difficult to harden and treat, further enamel hardening occurs within a short time after eruption in pigs, allowing for healthy tooth function (Biondi et al. 2017; Depalle et al. 2023). This requires a posteruptive maturation where mineral density increases while protein content decreases. Proteins, such as albumin, that bind to enamel crystals and prevent crystal growth and enamel hardening have been suggested as cause for hypomineralized human enamel that does not naturally harden after eruption. However, albumin is abundant in pig enamel. It is unclear whether fast posteruptive enamel hardening in pigs occurs despite the high protein content or requires a facilitated protein loss to allow for crystal growth.

After eruption, enamel is immersed in saliva that is rich in enzymes and ion. Posteruptive maturation mechanisms that occur over many weeks and years through the diffusion of calcium and phosphate from saliva were previously suggested (Lynch 2013). However, it remains unclear how the rapid mineral density increase in pigs occurs despite the high protein content in porcine enamel (Gil-Bona et al. 2023). Previous studies have shown that pig teeth have a remarkably low mineral content and high content of albumin (Kirkham et al. 1988; Robinson 2014). In addition, serum albumin has been shown to bind to enamel crystals, herby arresting crystal growth (Robinson et al. 1998; Shore et al.2000). The desorption of albumin and its removal from the crystal surface are required for extended enamel crystal growth and to attain fully mature enamel. The main enzymes responsible for protein removal during enamel development, MMP-20 and KLK-4, are secreted by ameloblasts before eruption and have been shown ineffective against the non-enamel serum proteins such as albumin (Bartlett 2013; Williams et al. 2020). Ameloblasts undergo apoptosis prior to tooth eruption and erupted enamel is cell-free. Therefore, an alternative protein removal is necessary to complete the enamel maturation after eruption in pigs to increase mineral density and achieve the required hardness.

We asked here how posteruptive changes in the enamel proteome track with mineral accumulation to better understand how porcine enamel hardens through diffusion-driven processes in the absence of cell-mediated transport. The objective of this study was to unravel the changes in the organic phase of enamel after tooth eruption and identify enzymes potentially involved in the posteruptive maturation process. To this end, we analyzed the enamel proteome from same tooth type at the time of eruption (PE-0), 2 weeks (PE-2), and 6 weeks (PE-6) after eruption, expanding on known mineral changes at these three post-eruptive time points (Depalle et al. 2023). Additionally, we analyzed the saliva proteome of the same animals at these time points to investigate potential influx or exchange between enamel and saliva.

The findings from this study will advance our understanding of posteruptive, diffusion driven enamel hardening by revealing the temporal changes in enamel proteome, endogenous and saliva-derived proteins in enamel, and potential exchange with saliva.

## MATERIALS & METHODS

### Animals

Yorkshire pigs at the age of 2-, 4- and 8-weeks-old (N=3 per age group) were obtained from Tufts University Cummings School of Veterinary Medicine following approved IACUC regulations (protocol number G2019-72). Animals were weaned after the age of 4 weeks, did not receive any antibiotics, and were sacrificed using intravenous injection of phenytoin pentobarbital sodium (Euthosol, Virbac, France) immediately followed by exsanguination to reduce bleeding and contamination of teeth during dissection.

### Saliva Collection

Using a saliva collection kit (SuperSAL601, Oasis Diagnostic Corporation, Vancouver, WA) approximately 1 mL of saliva was obtained prior to sacrifice. Saliva samples were immediately transferred onto dry ice upon collection and stored at -80°C until being processed. Samples were centrifuged at 14,000 x g for 10 minutes at 4°C, and the supernatant was concentrated using a 10,000 MW filter unit (Vivaspin 500, Vivaproducts, Littleton, MA) at 12,000 x g. The retentate was washed twice in extraction buffer (8 M urea, 50 mM Tris pH 8.0, 10 mM DTT) and the proteins were resuspended in 50 mM iodoacetamide (IAA) and 50 mM ABC buffer. After 15 minutes the mixture was centrifuged at 12,000 x g for 15 minutes. The retentate was washed and resuspended in 50mM ABC buffer and stored at -20°C until gel electrophoresis (Walz et al. 2009)

### Enamel Collection

After careful dissection without any blood contamination, enamel surface was further cleaned by brushing with phosphate buffered saline. Three quarters of the erupted enamel thickness was collected from the same tooth type in all animals (primary 4^th^ mandibular premolars) using carbide burs on a low-speed drill. For each timepoint three teeth were used, one tooth per animal. Disposable items were used for each sample when possible; surfaces and non-disposable items were cleaned with 70% ethanol between each sample.

### Protein extraction from enamel

was done according to our previously published work (Green et al. 2019). 10 mg of enamel powder from each sample was incubated with 12% TCA for 48 hours at 4°C under agitation. Samples were then centrifuged at 4°C for 45 min (2,500 x g) and washed twice using acetone. The resulting pellet was resuspended in a suspension buffer (3M urea + 50 mM ammonium bicarbonate buffer [ABC]) and stored at -20°C until gel electrophoresis

### Protein precipitates from enamel and saliva samples

were loaded into gels (BioRad) and were subjected to in-gel trypsin digestion (Shevchenko et al. 1996; Gil-Bona et al. 2023). Peptides were subjected to electrospray ionization and then entered into an LTQ Orbitrap Velos Pro ion-trap mass spectrometer. Peptide sequences (and hence protein identity) were determined by matching protein databases with the acquired fragmentation pattern by the software program, Sequest (Thermo Fisher Scientific, Waltham, MA) (Eng et al. 1994). All databases include a reversed version of all the sequences and the data was filtered to between a one and two percent peptide false discovery rate. Proteins with only 2 or more peptides were included in the analysis.

The analysis of the proteomics counts data matrix was implemented with the package edger (Robinson et al. 2010). Cluster analysis plots were generated with the t-sne implementation from the “rtsne” package, the principal coordinates function (pco(…)) from the “labdsv” package and the base pca (prcomp(…)) function. Volcano plots were generated with the base plot (plot(…)) function and peptide heatmaps were generated with the heatmap (heatmap.2(…)) function from the “gplots” package. Other r packages used were “corrplot” to visualize correlations across the sample matrix, “openxlsx” to read and write excel compatible files, “ggplot2” for plotting, and “venndiagram” for plotting venn diagrams. Other figures and heatmaps were generated using Prism 10 (Graphpad Software Inc., San Diego, CA, USA) and relative abundancies in the heatmaps are based on the intensity and area of the most abundant peptides.

## RESULTS

### 2.1. Proteins in enamel decrease within two weeks after eruption

We identified 1035, 456 and 250 proteins in enamel at eruption (PE-0), two weeks after eruption (PE-2) and six weeks after eruption (PE-6), respectively (Figure 1A). 597 of those proteins were only identified at PE-0, while 61 and 24 of them were unique to the PE-2 and PE-6 enamel respectively. The number and abundance of proteins decreases six weeks after eruption but most dramatically within two weeks after tooth eruption, between PE-0 and PE2, (SI Fig. 1)

**Figure 1.**
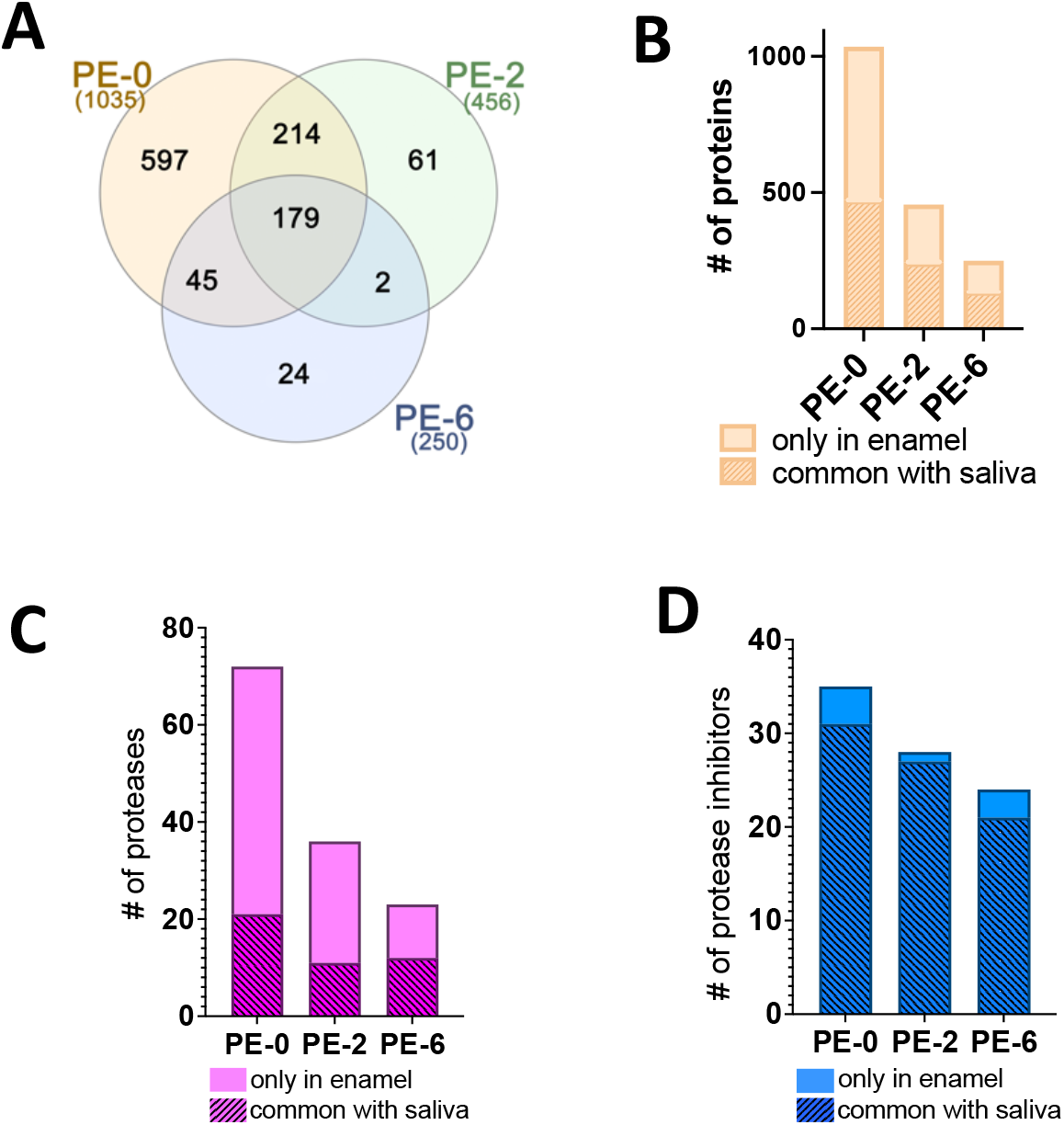
**A)** Number of proteins identified at each timepoint in enamel samples, and the number of unique proteins for each timepoint. **B)** Number of proteins identified in enamel decreases with posteruptive time and nearly half those proteins are common with saliva of the same animals. **C)** Number of proteases found only in enamel deceases drastically whereas the number of those shared with saliva change little with time. **D)** Protease inhibitors found only in enamel and those shared with saliva decrease similarly with posteruptive time.

**Figure 2.**
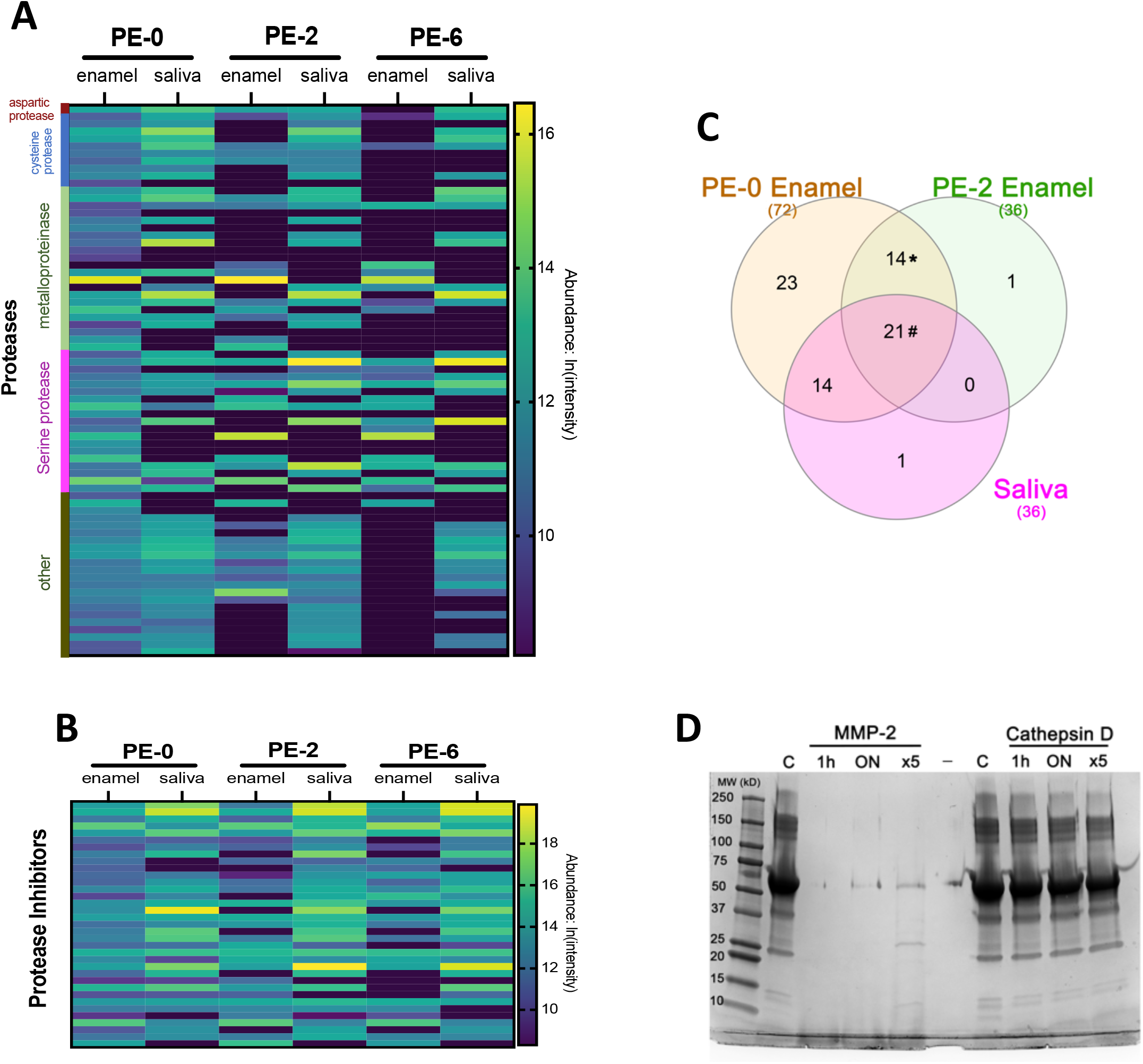
The relative abundancies of the **A)** proteases and **B)** protease inhibitors identified in enamel and saliva seen in heatmaps with identified proteases grouped into the cysteine, aspartic, serine and metalloproteinase protease families. In enamel, both diversity and abundance of proteases decrease with time after eruption. Abundancies and the diversity of the protease inhibitors in enamel remain relatively unchanged throughout the posteruptive time, when compared with the temporal change in proteases. **C)** Venn diagram of proteases detected in enamel only at PE-0 and PE-2 and proteases seen in saliva at all time points. The list of proteases corresponding to the fields of this Venn diagram can be found in Table 1. **D)** Activity of two enzymes that persist in enamel against porcine serum albumin.MMP-2, one of the enamel proteases marked by the asterisk in the Venn diagram, cleaves albumin rapidly at 37°C. Cathepsin D, found in the porcine saliva at all ages and one of the proteases marked by the number sign in the Venn diagram, did not show activity against albumin even at higher concentrations.

### 2.2 Saliva infiltrates enamel during posteruptive maturation

The number of proteins found in enamel that are in common with saliva constitutes half of the total proteins in enamel at all three timepoints, 45.7% at PE-0, 53.2% at PE-2 and 55.2% at PE-6 (Figure 1B), as both enamel and saliva proteins decrease in abundance with time. In addition, the number of proteases in enamel also decreases with time, from 72 at PE-0, to 36 at PE-2 and 23 at PE-6 (Figure 1C). Number of protease inhibitors in enamel also follows the same trend with, 35 at PE-0, 28 at PE-2 and 24 at PE-6 (Figure 1D). Unlike the gross number of proteins in enamel, the number or proteases and protease inhibitors that are in common with the saliva of the same animals stay relatively stable throughout the 6 weeks of posteruptive time and do not follow a stable ratio described in Figure 1B. 21, 11 and 12 of the proteases were in common with saliva and 31, 27 and 21 of the protease inhibitors were in common with saliva at PE-0, PE-2 and PE-6 respectively.

### 2.3. Enamel Specific Proteins and Albumin Decrease in Abundancy After Eruption

The total number of unique peptides from enamel proteins in porcine enamel decrease with posteruptive time (Figure 3A), while the number of peptides from KLK-4 stays at 3 peptides in each timepoint. However, the high levels of MMP-20 follow the trend of decrease with posteruptive time, 22 in PE-0, 18 in PE-2 and 11 in PE-6 (Figure 3B). Unique peptides representing serum albumin also decreases, 26 in PE-0, 23 in PE-2 and 20 in PE-6 however peptides from the fetal isoform stays relatively stable with 11 at PE-0 and 9 at PE-6 (Figure 3C). The abundancies of each protein based on the area of the most abundant peptides shows similar trends with the peptide numbers and can be seen in the heatmap in In Figure 3.D. Adding to our previous findings, we confirmed the presence of serum albumin in enamel samples using ELISA (SI sheet 1). As indicated inside the cells of the heatmap in Figure3D, the percentage of the amino acid sequence covered by the identified peptides for each protein also decreases with posteruptive time.

**Figure 3.**
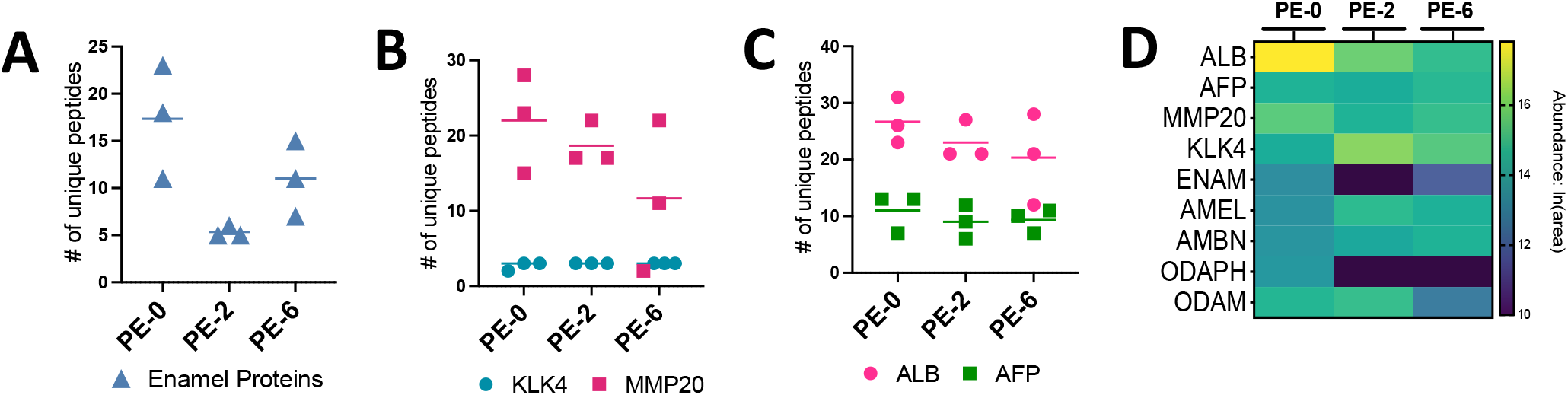
The number of unique peptides identified at each timepoint for **A)** Enamel proteins (AMEL, ENAM, AMBN, ODAM, and ODAPH combined) **B)** Enamel proteases KLK4 and MMP20, **C)** Serum albumin (ALB) and its fetal isoform (AFP) **D)** Heatmap showing relative abundancies of these proteins based on the area of the most abundant three peptides.

### 2.4. GO Analysis Reveal Enrichment of Protease Inhibitors after Eruption

A categorization of the enamel proteome at the three posteruptive timepoints, based on Biological Processes, Molecular Function, and Protein classes, using the Panther database (Mi et al. 2019) can be seen in Figure 4,A-C. All enamel samples had the highest number of proteins in the class of binding and catalytic activity of Molecular Function. Similarly, all enamel samples had the highest number of proteins for biological processes in the classes of cellular and metabolic processes. The only difference between PE-0, PE-2, and PE-6 was in the protein class category (Figure 4, pink bars). At all timepoints, the enzyme class of metabolite interconvention was dominant, which includes the class of oxidoreductases. However, in PE-0 enamel the class of protein modifying enzymes (protease class) represents the second highest percentage of proteases, while at PE-6 the second highest percentage of proteases pertains to the class of protein binding activity modulators, which includes protease inhibitor class (Figure 4A, B, C). The number of both proteases and protease inhibitors decreases from PE-0 to PE-6 (Figure 1 C,D), however in the GO enrichment analysis (Ashburner et al. 2000), the fold change of the protease inhibitors changes from 5 to 15 between PE-0 and PE-6, while the protease fold change remain stable (Figure 4D).

**Figure 4.**
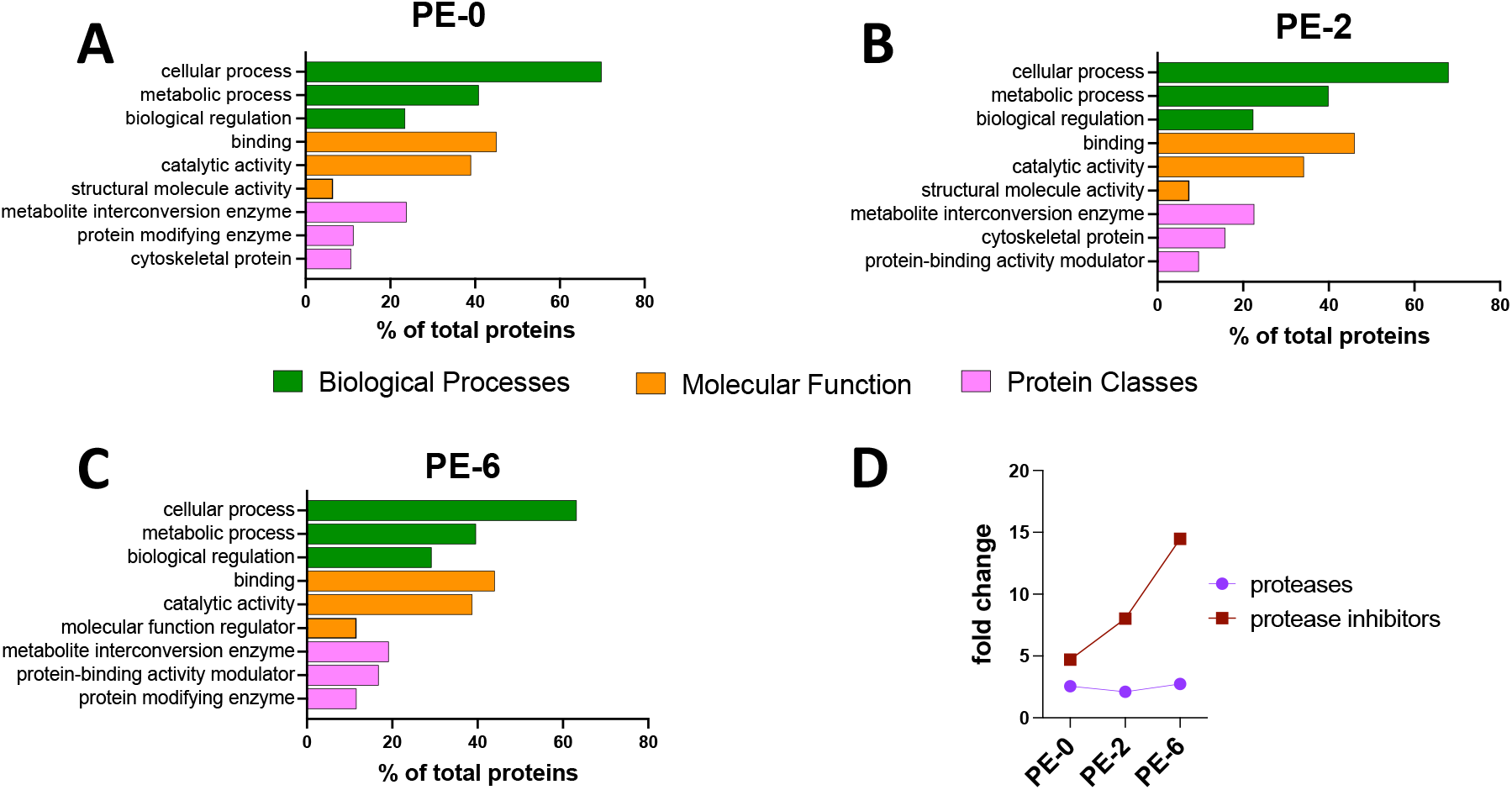
**A, B, C)** Top three families of enamel proteins under the categories of biological processes, molecular function, and protein classes from PantherDB for each timepoint. **D)** Based on the GO Enrichment analysis on normalized number of proteins, the fold change of proteases remains stable at 2, while the protease inhibitors have an increasing fold change, from 5 (PE-0) to 15 (PE-6).

## DISCUSSION

Posteruptive increase in enamel mineral density and changes in crystal morphology over time have been reported for human enamel previously (Wöltgens et al. 1981; Driessens et al. 1985; Kotsanos and Darling 1991; Wang et al. 2005; Kataoka et al. 2007; Lynch 2013; Wada et al. 2023). In these studies, posteruptive enamel maturation has been described as a passive and slow process, that occurs through deposition of ions and mineral delivered by saliva (Driessens et al. 1985; Palti et al. 2008). In contrast to the relatively slow development of human teeth, the very fast development of pigs and pig teeth results in teeth that erupt soft but quickly need to meet the biological requirements of mastication for an omnivorous diet. This necessitates a mechanism of posteruptive tooth enamel hardening that is both fast and effective despite the high protein content in pig enamel at the time of tooth eruption.

The incomplete protein removal in pig enamel before eruption and the correlation between the protein abundancy and mineral accumulation during enamel maturation has been reported previously (Robinson et al. 1988). Detailed information on the time course of posteruptive changes in the proteome of pig enamel has been lacking. This study addressed this gap in knowledge, and our data show how protein diversity and abundancy decrease quicky in porcine enamel after eruption when teeth are exposed to the oral environment and enamel is in contact with saliva. Our comparison between the proteome of enamel at the time of eruption, and 2 and 6 weeks after eruption, showed a dramatic decrease of protein content in enamel within two weeks after eruption. Although not as drastic, the protein content continues to decrease until 6 weeks after eruption. These data complement and are in line with our previous findings of a rapid increase in mineral density right after tooth eruption. In a crystallographic study, Kallistová and colleagues also showed that the final enamel maturation in pigs is obtained shortly after eruption (Kallistová et al. 2018).

A saliva-guided mechanism that finalizes enamel maturation after tooth eruption in pigs has been suggested in the past (Kirkham et al. 1988; Robinson 2014). Our earlier work has shown that the surface enamel in pig teeth is very porous at eruption and gains quickly in mineral density within just weeks after eruption (Depalle et al. 2023).Despite the high porosity of pig enamel, major saliva-specific proteins, such as mucin, statherin, or histatin protein families, were not detected in the porous enamel. However, the important role of saliva in posteruptive maturation becomes apparent in the number of proteins that are detected in enamel and shared between saliva and enamel. The number of proteins that are shared in enamel and saliva amounts to about half of the number of total proteins in enamel and this ratio remains stable at all three timepoints from eruption to 6 weeks post eruption.Nevertheless, certain classes of proteins, such as proteases and protease inhibitors change in relative abundance with time and the portion of individual protease subclasses (serine, cysteine) at each timepoint. The changing abundance of saliva derived proteins and proteases detected in enamel is likely related to enamel permeability and porosity, electrical charge of the proteins, pellicle composition, and possibly the plaque composition on the enamel surface (Zimmerman et al. 2013).

It has been shown for human hypomineralized lesions that deproteinization increases the success of current approaches for hardening and restoration (Ekambaram and Yiu 2016; Ekambaram et al. 2017; Sönmez and Saat 2017; Fernando et al. 2020). One challenge for successful mineralization treatments and restorations is that the unique protein composition of the demarcated opacities, specifically the abundance of albumin, impedes enamel crystal growth and needs to be considered.Deproteinization using sodium hypochlorite likely degrade the organic phase in enamel without discrimination and might compromise the mineralization and healthy physical properties of enamel that benefit from small quantities of retained organic matter in mature enamel (Gibson 2011).

The effective posteruptive mineralization in pig enamel requires proteolytic activity to degrade serum albumin in addition to other proteins trapped in hypomineralized enamel. Enzymes to facilitate such protein removal have to be present in at least both PE-0 and PE-2 enamel and, if derived from saliva, should be present for weeks after the different teeth within the dentition erupt (Table 1). While the MEROPS peptide database is useful for predicting substrate specificity of proteolytic enzymes, this database is not complete and might miss potential candidates for posteruptive proteases in enamel (Rawlings et al. 2014). In addition, physiological conditions and tissue specific properties might affect the activities of listed proteases against albumin and other proteins (Kragh-Hansen 2018). For instance, reports on albumin precursor and albumin as a substrate for KLK4 or MMP20 are conflicting (Matsumura et al. 2005; Williams et al. 2020). In addition, serum albumin has been reported as a potential substrate for other MMPs (MMP2) and members of the cathepsin family (Schilling and Overall 2008). We tested the albuminase activity of enzymes from these two families, MMP2 and Cathepsin D, which are also identified in our samples (Figure 2D). While MMP2 was able to degrade albumin rapidly, the effect of Cathepsin D against albumin was insignificant.

**Table 1.** List of candidate enzymes responsible for the posteruptive maturation of porcine enamel. Based on the proteases shown on Figure 2A, we listed the proteases that are present at least in both PE-0 and PE-2 enamel. Additionally, any salivary protease needs to be identified also in the saliva at all ages. The “pseudoenzymes” and the coagulation factors are not listed in this table.

| Protease Name† | Protease Family | Also Found in Saliva?* | Ameloblast expression | Activity Against Albumin | Activity Against Enamel Matrix Proteins | Reference |
| --- | --- | --- | --- | --- | --- | --- |
| Cathepsin D* | Aspartic | (+) | (+) | (-) * | N / A | (Tye et al. 2009; Chun et al. 2024) |
| Acid Ceramidase | Cysteine | (+) | N / A | N / A | N / A | - |
| Cathepsin B | Cysteine | (+) | (+) | (+) | (+) | (Smid et al. 2001; Tye et al. 2009; Lee et al. 2014) |
| Cathepsin C (dpp-1) | Cysteine | (-) | (+) | N / A | N / A | (Tye et al. 2009) |
| Cathepsin L | Cysteine | (-) | (+) | N / A | N / A | Tye 2008 |
| Carnosine Dipeptidase 2 | Metalloproteinase | (+) | N / A | N / A | N / A | - |
| Peptidase D | Metalloproteinase | (+) | (+) | N / A | N / A | (Simmer et al. 2014) |
| MMP-20 | Metalloproteinase | (-) | (+) | (-) | (+) | (Lu et al. 2008; Williams et al. 2020) |
| MMP-2* | Metalloproteinase | (-) | (+) | (+)* | (+) | (Caron et al. 2001) |
| Dipeptidyl-peptidase 2 (dpp-7) | Serine | (+) | (+) | N / A | (+) | (Smid et al. 2001; Maes et al. 2007; Tye et al. 2009) |
| Elastase | Serine | (+) | N / A | N / A | N / A | - |
| Esterase D | Serine | (+) | N / A | N / A | N / A | - |
| KLK-4 | Serine | (-) | (+) | (-) | (+) | (Lu et al. 2008; Williams et al. 2020) |
| Carboxypeptidase E | Other | (-) | (+) | N / A | N / A | (Simmer et al. 2014) |
| Proteasome Subunits A, B ,C | Other | (+) | (+) | (+) | (+) | (Sixt et al. 2007; Geng et al. 2015) |

\* Based on the data from this study.
†Proteases that are found in PE-0 and PE-2 enamel.
N / A: Data not available

In addition to MMP2, MMP20, the main enamel protease expressed in the secretory stage of enamel development (Caron et al. 2001), was found in high levels in the porcine enamel, consistent with a continuous role of residual MMP20 during posteruptive maturation enamel (Bartlett 2013). The other major enamel protease with a key role for enamel maturation is KLK4, which has also predicted cleavage sites in serum albumin (Rawlings et al. 2014), although it has been shown ineffective degrading albumin (Williams et al 2020).

Posteruptive enamel maturation include interactions between enamel crystallites and the organic phase of the enamel, as well as processes of protein removal facilitated by residual enamel enzymes, salivary proteases, and exchange between enamel and saliva (Goodson et al. 2017). A highly porous surface of recently erupted porcine enamel allows these processes to promote a decrease in protein content and allow for further mineral accumulation. This study shows that enamel composition changes quickly after eruption and that the abundance of protein continues to decrease as the tooth spends more time in the oral cavity. Further *in vitro* and *ex vivo* studies are needed to fully understand the physiological mechanisms of post-eruptive enamel maturation.

## Supporting information

SI sheet 1

## ACKNOWLEDGEMENTS

This work was supported by NIH grants R01DE025865(FBB), R21DE029903-01(FBB) and R90DE027638(HK). Parts of this work were performed at the Taplin Mass Spectrometry Facility, Cell Biology Department, Harvard Medical School.

## Notes

### Competing Interest Statement

The authors have declared no competing interest.

### Summary of Updates

Included the acknowledgments section with funding information.

