## Supplementary figures and images for "Posteruptive Loss of Enamel Proteins Concurs with Gain in Enamel Hardness"

### SI Fig.1

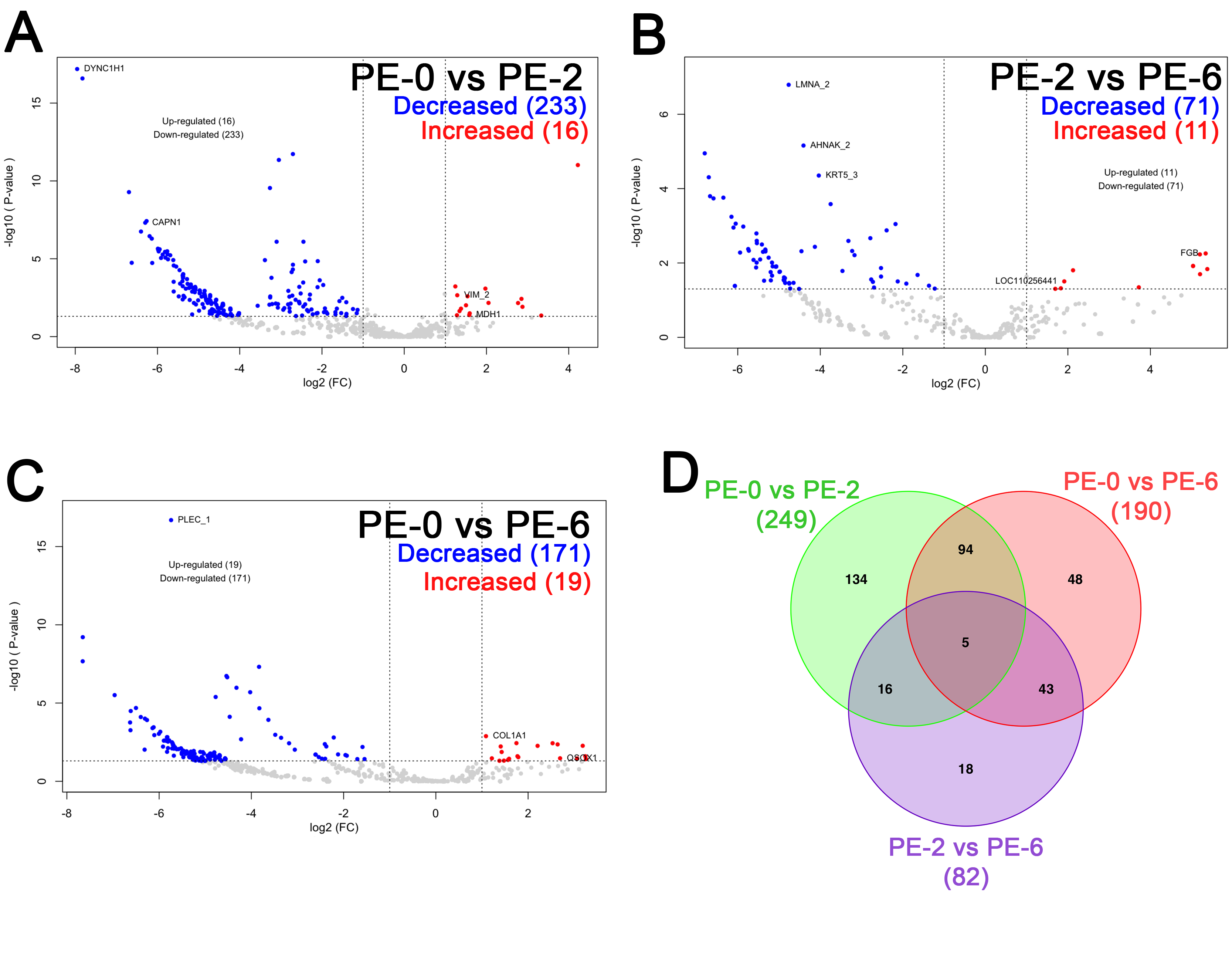
