## Supplementary material for "Posteruptive Loss of Enamel Proteins Concurs with Gain in Enamel Hardness": SI sheet 1

| | conc | dil ng/mL | ng/ul | ng albumin in 10mg pig enamel (25ul sam |
|---|---|---|---|---|
| en1 | 8.4825 | 373.23 | 0.37323 | 9.33075 |
| en2 | 12.1405 | 445.151667 | 0.44515167 | 11.1287917 |
| en3 | 13.0025 | 476.758333 | 0.47675833 | 11.9189583 |

ıple vol )

| | |
|---|---|
| | **in 10mg enamel** |
| **albumin (ng)** | **10.7928333** |
| **std** | **1.3264074** |
